# Identifying key federal, state and private lands strategies for achieving 30x30 in the US

**DOI:** 10.1101/2021.03.26.437234

**Authors:** Lindsay M. Dreiss, Jacob W. Malcom

## Abstract

Achieving ambitious conservation goals to conserve at least 30% of US lands and waters by 2030 (“30x30”) will require a multi-scale baseline understanding of current protections, key decision makers, and policy tools for moving forward. To help conservationists and decision makers support the science-based call to address the biodiversity and climate crises, we analyze the current spatial patterns of imperiled species biodiversity and carbon stores in the U.S. relative to protected areas. Analyses demonstrate that 30x30 is numerically achievable nationally, but high spatial heterogeneity highlights the need for tailored approaches from a mix of authorities at federal, regional, and state scales. Critically, current land protections rarely overlap with areas essential for conserving imperiled species biodiversity and mitigating climate change. Nearly one-fifth of unprotected biodiversity hotspots and carbon-rich areas are also at risk of either land conversion or climate exposure by 2050. We discuss this baseline relative to key policy considerations for making practical, substantive progress toward the goal.

## Introduction

According to the international conservation community, goals for conserving biodiversity cannot be met given current trajectories of environmental degradation and without transformative changes across economic, social, political, and technological spheres (1). Restoration and maintenance of quality habitat through a more extensive global protected areas (PAs) network (see Aichi Target 11; 2) is considered essential for conservation. Yet the global community has fallen short of 2020 targets for PA coverage at a time when threats to biodiversity – foremost, habitat conversion – are at an all-time high (3).

Recognizing the critical role that PAs have in conserving biodiversity and mitigating climate change impacts has led to increased interest in adopting new national and international conservation targets. The Global Deal for Nature, a science-driven plan to sustain biodiversity and address climate change, calls for at least 30% of Earth to be formally protected by 2030 (“30x30”; 4). The Convention on Biological Diversity has drafted a post-2020 global framework that includes 30x30 as a steppingstone toward a 2050 Vision for Biodiversity (5). In the United States, the number of proposed policy measures aligning with a 30x30 framework is on the rise. As of early 2021, this includes a federal executive order and a California state executive order (6, 7). These efforts provide an opportunity to integrate biodiversity and climate agendas and promote land protections that can maximize biodiversity conservation and minimize carbon loss at multiple scales.

Past research on setting PA priorities for biodiversity (8,9,10) and natural climate strategies (11,12) provide a useful foundation for addressing this need, but do not align with the policy tools and units at which federal and state decision-makers govern. To achieve 30x30 in a way that most benefits biodiversity and climate, policy makers need guidance on how to operationalize these targets.

Here we synthesize biodiversity and ecosystem carbon data with policy-relevant land protections at multiple scales to provide a baseline assessment for charting a path to 30x30 in the U.S. We use the Protected Areas Database of the U.S. (13) to spatially define areas that are currently managed for biodiversity conservation (GAP 1 & 2) and that may be in need of additional protections (GAP 3 & 4) (Panel 1). We compare GAP classified lands with lands rich in imperiled species and carbon to estimate how much of these areas are already protected and where new protections would yield larger gains for biodiversity conservation and climate mitigation objectives. In addition, we assess the risk of these areas of being lost due to conversion or threatened by advances in climate change. These results are not meant to serve as a map of priority lands, but as a basis for framing a discussion on operationalizing the key objectives of 30x30. Our aim is to help conservationists and decision-makers plan and take critical next steps to operationalize 30x30-in doing so, addressing some of the greatest conservation challenges faced by the United States and the world.

## Results

Twelve percent of lands within the U.S. and its territories are generally managed consistently with biodiversity conservation goals (i.e., GAP 1 and 2). Up to 29.8% of U.S. lands and territories are managed for either biodiversity conservation or multiple uses (i.e., GAP 1, 2, and 3). This leaves a large majority of the U.S. lacking any known protections from land conversion (70.2%; i.e., GAP 4).

Areas managed for biodiversity (85.7% of GAP 1 and 2 areas) and for multiple uses (85.6% of GAP 3 lands; Figure 1a) fall largely on federally managed lands. Protecting 30% of lands could nearly be achieved at the national scale if new conservation-based mandates were applied to all federally managed GAP 1-3 areas (27.7%).

**Fig. 1.**
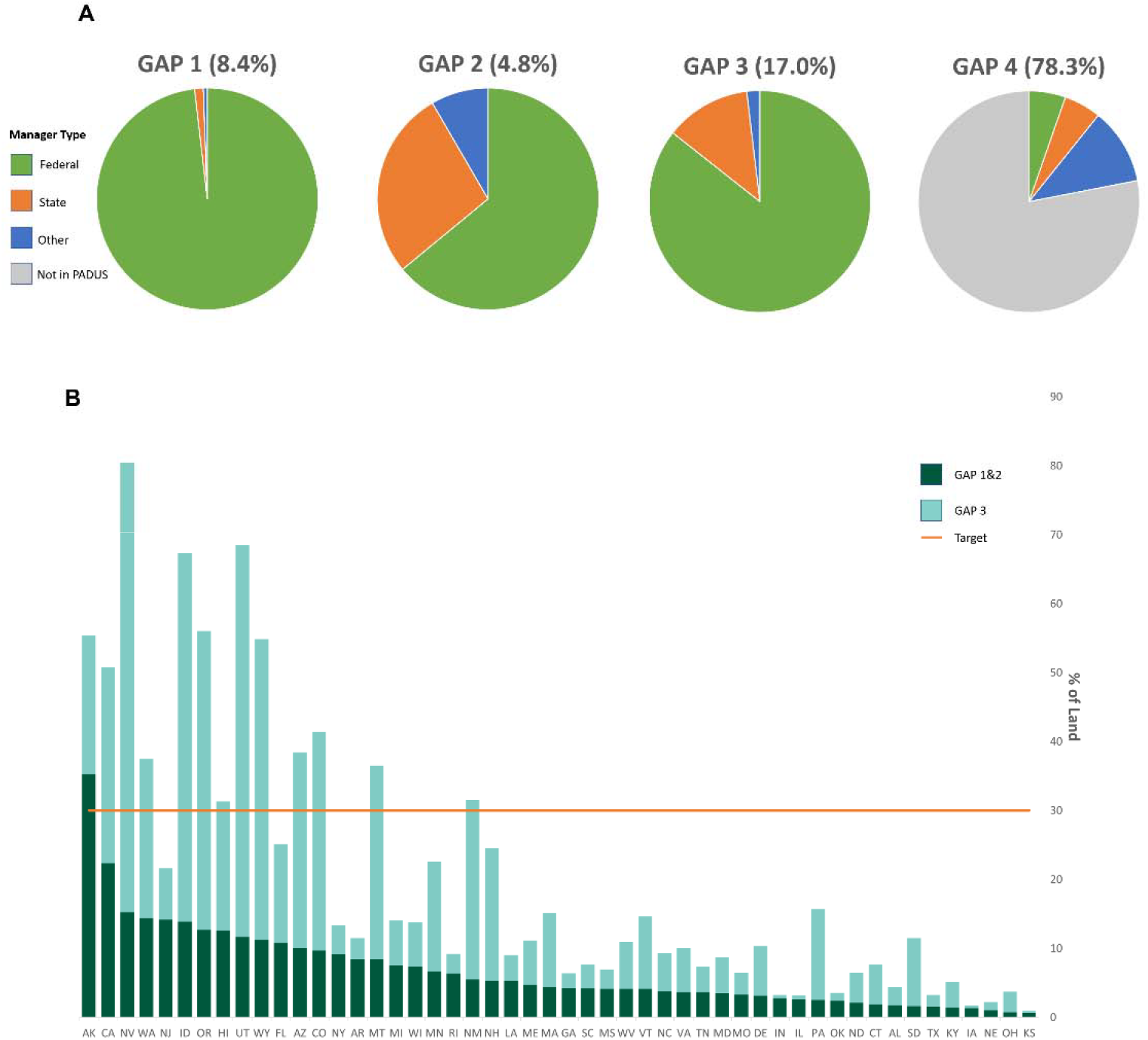
Proportion of U.S. lands in PADUS by GAP status code, manager type and state. (A) federal agencies manage much of the protected areas network and (B) most states have few protections for biodiversity as they fall well-short of the 30% target given existing and potential (GAP 1-3) protections.

At the state-level, Alaska is the only state to have at least 30% of its territory managed for conservation (Figure 1b). Twelve western states - California, Nevada, Washington, Idaho, Oregon, Hawaii, Utah, Wyoming, Arizona, Colorado, Montana and New Mexico - would achieve the 30% numerical target if GAP 3 areas were similarly managed. Generally, federal agencies manage the majority of GAP 3 lands in these states. In contrast, the 11 states with greater state-level authority are located in the Northwest and Midwest (Table S1).

Imperiled species biodiversity hotspots and GAP 1 or 2 areas rarely overlap, with only 7% of hotspots covered by GAP 1 and 2 lands (Figure 2). Similarly, 12.8% of highest potential areas fall within GAP 1 and 2 lands. While including GAP 3 lands significantly increases the coverage for biodiversity hotspots (from 7% to 20%; Figure 2), 80% of the most diverse areas would still remain unprotected because they fall on GAP 4 or otherwise unprotected private lands. Nearly two thirds of the top carbon-rich areas fall in GAP 4 areas. While rarity-weighted richness exhibits similar broad spatial patterns to raw species richness (Figures 1 and 2), there is significantly greater overlap with key GAP lands: 32.6% of rarity hotspots are covered by GAP 1 and 2 areas and an additional 20.5% by GAP 3 lands.

**Fig. 2.**
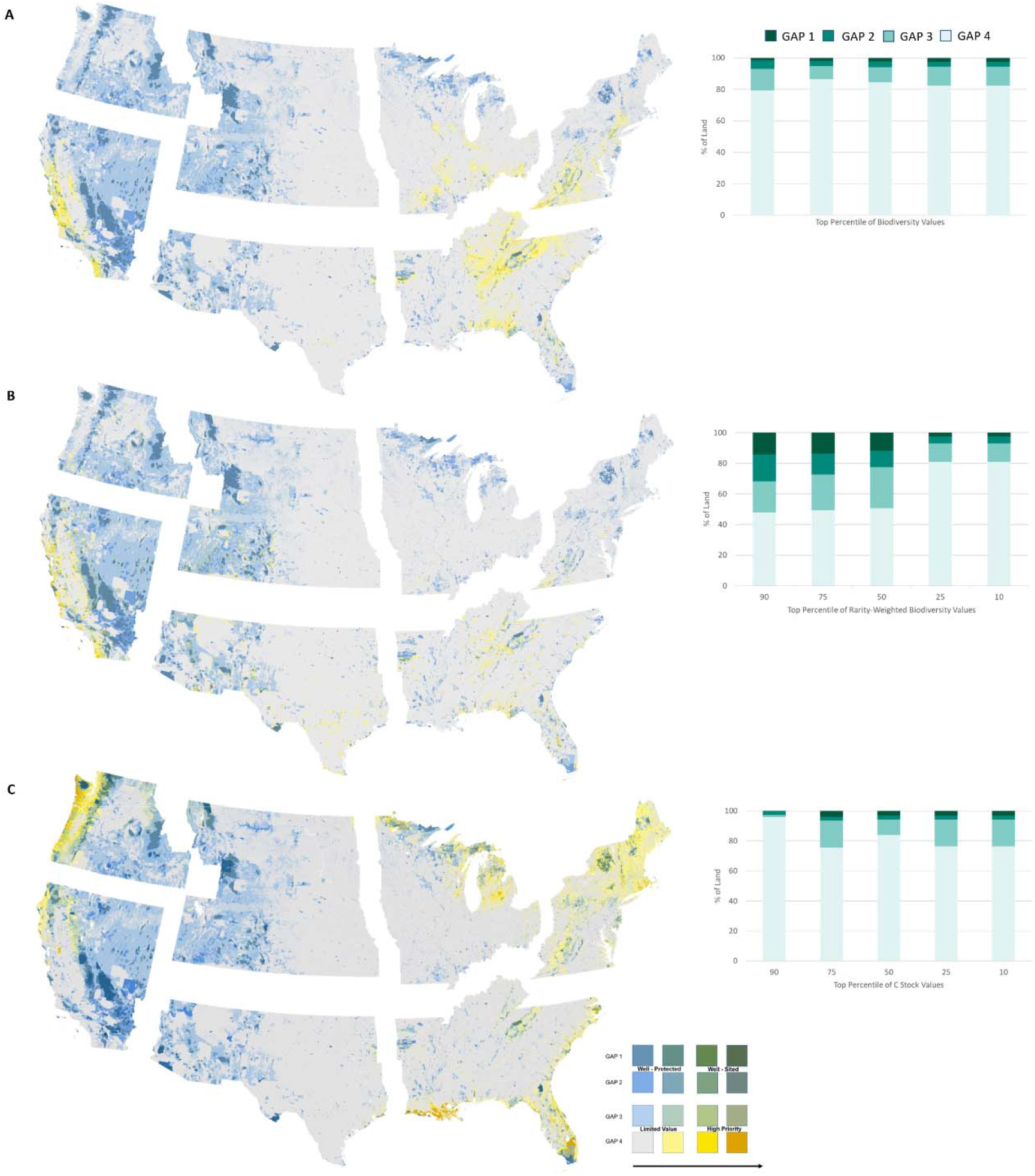
Combining protected area coverage to locations of biodiversity hotspots and carbon-rich areas. A) imperiled species richness, B) rarity-weighted richness and C) ecosystem carbon (tonnes C per hectare) show significant areas of mismatch that need to be addressed. The blue (y-axis) component of the bivariate color ramp signifies protections under GAP categories 1-4 while the yellow (x-axis) component signifies the corresponding 30x30 objective based on quantile intervals where the top 10% of values are in yellows (imperiled species richness or ecosystem carbon). Bar charts represent the overlaps of different percentiles of biodiversity or carbon values with GAP categories (green color ramp). Biodiversity and carbon hotspots were considered as locations with values in the top 90^th^ percentile of the distribution.

As expected, the goals of protecting areas of high biodiversity and areas of high carbon mitigation potential are not completely decoupled: 22.3% of the top quantile of species richness locations (25.6% of rarity-weighted richness hotspots) are also very carbon-rich. These percentages nearly double when assessing overlap with the top 30% of carbon-rich areas. Unfortunately, few areas that meet both needs are managed in ways to help sustain them: 17.4% of areas that serve as hotspots for biodiversity and carbon fall on GAP 1 or 2 lands (<0.01% of CONUS).

### A 30x30 Typology

To guide operationalization of the 30x30 framework, we use GAP categories as proxies for policy options and biodiversity and ecosystem carbon as representative of the underlying goals of 30x30 to describe four basic types of classifications (Figure 2 legend):

#### Well-sited

Areas with long-term conservation management mandates (GAP 1-2 coverage) and high biodiversity and/or ecosystem carbon, where areas are effectively placed to achieve greater biodiversity conservation and climate mitigation. Priority actions for well-sited areas include maintaining existing protections and expanding protections outward in a way that expands landscape connectivity.

#### High Priority

Areas of overlap between weaker or short-term mandates (GAP 3-4) and greater biodiversity and/or ecosystem carbon, where new PAs or more protective policies would do the most to protect biodiversity and mitigate climate. These are the areas of greatest opportunity, where efforts to expand PAs would have especially high returns on investment.

#### Well-protected

Areas with long-term conservation mandates but relatively low local imperiled species diversity and/or ecosystem carbon potential, indicating lower return on conservation goals than other areas might provide. Despite their lower biodiversity and carbon stocks, these areas may serve as anchors for expanding protections to key adjacent areas.

#### Limited Value

Areas with weaker or short-term mandates and low levels of biodiversity and/or ecosystem carbon potential, indicating a low return on protections and an advantage to site PAs elsewhere. These are the lowest priority areas for 30x30. However, it is important to note that basing decisions solely on current habitat and species ranges may discount the necessity for future habitat recovery and restoration given climate change (14).

### Future Risk for Unprotected Hotspots

The level of risk faced is highly dependent on the type of risk and geographic location. Over 18.7% of biodiversity hotspots and 19.5% of carbon rich areas fall into the top category of at least one of the risks analyzed. Land conversion was the greater risk for biodiversity hotspots, with 20.8% of unprotected hotspots located in an area with at least 50% probability of conversion to non-natural land uses (Figure 3A). These at greatest risk of conversion were scattered along the Appalachian range, the Gulf Coast, Western California and areas of the Midwest. Fewer (8.2%) of carbon-rich areas faced the same probability of conversion, with these areas more concentrated in southern Florida, the Pacific Northwest, and Michigan (Figure 3B).

**Fig. 3.**
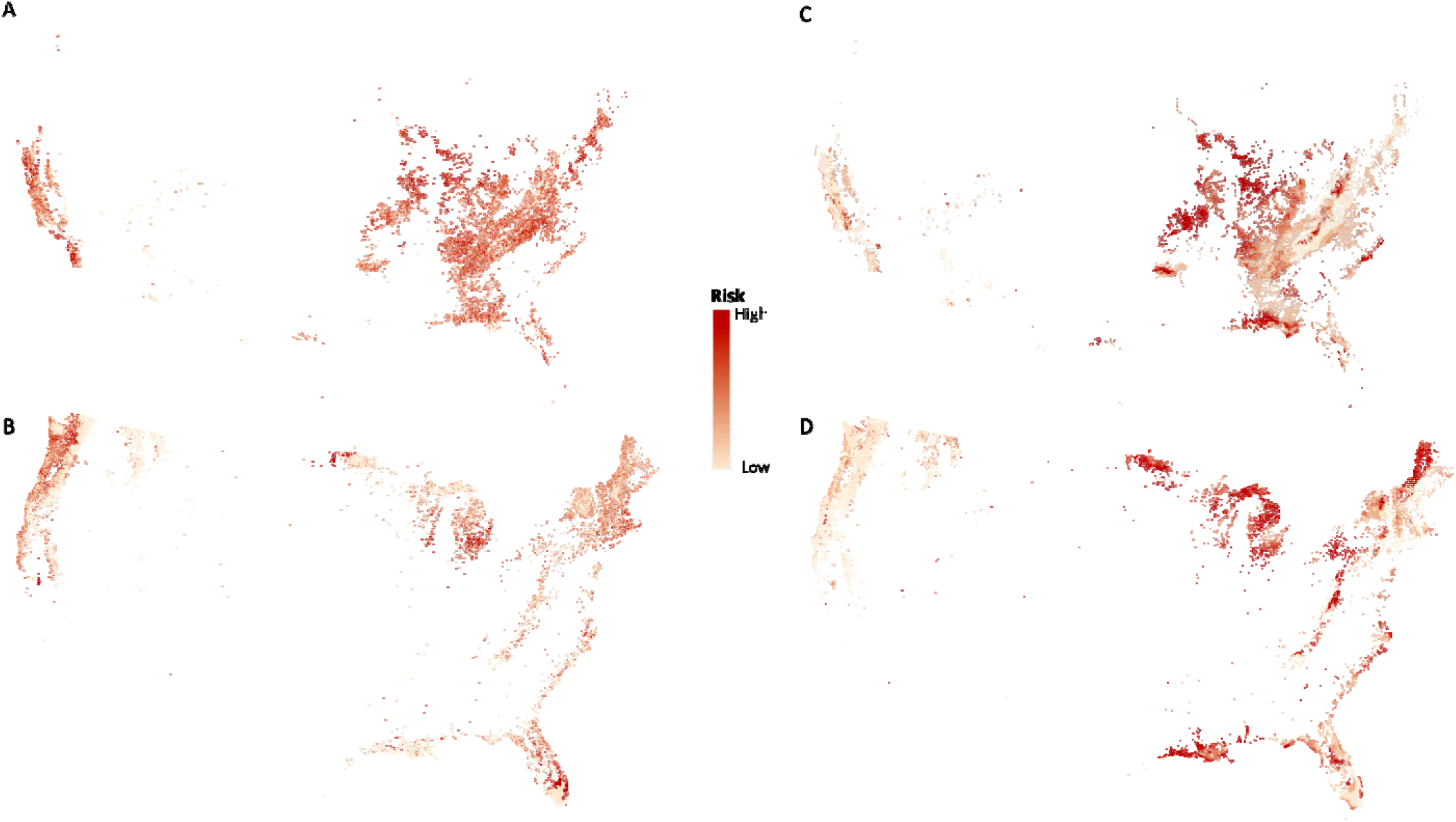
Levels of risk for unprotected biodiversity hotspots and carbon-rich areas. Unprotected areas face varying risk of conversion to non-natural land uses (A-B) and exposure to climate change (C-D) dependent on geographic location. Hotspots represent the top 90^th^ percentile of the distribution of values for species richness (A&C) and stored carbon (B&D). Risk is measured by probability of land conversion (on a scale of 0 – 1) and climate velocity (0 – 57 km/yr).

Unprotected biodiversity hotspots generally faced less exposure to climate change: 3.6% of areas overlapped with the top quantile of climate velocity values. These were located in the Ozark Mountains, Gulf Coast, and parts of the Midwest (Figure 3C). However, 15.6% of unprotected carbon-rich areas were subject to the highest climate exposures, particularly in Louisiana, upper Midwest, and New England (Figure 3D).

## Discussion

While the basic numerical accomplishment of protecting 30x30 is feasible at the national scale given the current extent of the PAs network, prioritizing biodiversity protection and climate change mitigation presents challenges and opportunities. In addition, high spatial variability in the distribution of PA designations and will require tailored approaches across regions (Figure 1).

An option for more rapidly reaching 30% includes establishing additional protections on GAP 3 lands. For example, nearly 30% can be achieved at the national level if regulatory changes to GAP 3 areas emphasized biodiversity protection over other uses. Because the majority of the PA network (GAP 1-3) is managed by federal agencies, action at the federal level may be essential to reach numerical goals. Land management laws like the National Forest Management Act and the Federal Lands Planning and Management Act afford species and habitats protections, but the effectiveness of protections may vary by ownership (15). Moreover, federal agencies can provide necessary leadership and coordination across jurisdictions to better ensure representation of all natural ecosystems (4).

Importantly, focusing on key federal lands alone would ignore the substantive goals of 30x30: 80% of biodiversity hotspots would still lack significant place-based protections if GAP 3 federal lands were converted. This result highlights the alarming reality that focusing strictly on the numerical goal of 30x30 could lead to outcomes contrary to intent, with new PAs being established in areas with low biodiversity or carbon mitigation potential (16). Regions with few public lands will face trade-offs between siting new areas based on biodiversity need or opportunity (e.g., isolated and sparsely populated; 17). Additionally, these areas will need to assess the future risks that come with failure to conserve biodiversity: nearly one fifth of all unprotected hotspots face high probability of land conversion and/or high potential climate exposures. GAP 4 areas are, by far, the most extensive, but would require more effort and investment from decision-makers to acquire land and/or establish biodiversity protections as priorities. A report to the National Climate Task Force on framing the implementation of 30x30 in the U.S. emphasizes the need for federal government to help local communities achieve their own conservation priorities and vision (18). Current federal conservation incentive programs, such as Farm Bill programs and those administered by the U.S. Fish and Wildlife Service are inadequate to address the need. As such, there must be significant efforts to advance conservation on private lands in key parts of the country.

In addition to federal and private lands, understanding state variation in biodiversity and protections is vital as important legal, social, and policy mechanisms operate at this level. We note twelve western states would achieve the 30% numerical target if GAP 3 mandates were strengthened. However, many areas of these states harbor few biodiversity or carbon-rich hotspots. In contrast, the 11 states where the state manages the majority of GAP 3 lands have higher biodiversity on average, but GAP 3 areas may not overlap with hotspots or be enough to significantly lessen the disparities in current PA coverage and the 30% target. State wildlife conservation programs employ state wildlife action plans that have the potential to advance conservation (e.g., 19), but are woefully underfunded (20). New state-level programs for public and private lands conservation can complement federal programs and create a more complete, multi-level solution.

Finally, we acknowledge the strong need for several additional considerations not accounted for in this work. In addition to the primary focus on biodiversity and climate, pursuing 30x30 will require addressing issues related to economic, political, and social constraints. For example, many High Priority areas for siting new protections also tend to be GAP 4 regions with higher human disturbance. This elevates the importance of restoration efforts (see, e.g., 21) and relative habitat condition could be integrated into analyses to avoid conserving areas with negligible conservation benefits. If properly planned, PAs in these areas may also provide opportunities for improving human health, well-being, and equitable access to nature. Goals to ensure a healthy environment for all communities have long been ignored or discounted in protected areas designations (22), in part because these topics are not well studied (23). Further research and planning are essential to ensuring access to quality nature for all.

This analysis is meant as a tool to aid decision-makers and not as a fully comprehensive plan. Our analyses are national in scope and intended to identify broad patterns to frame the national discussion; as such, local and domain-specific details are likely to vary.

Additionally, focusing on values at the national scale means that entire ecosystems important to representing local species assemblages and key ecosystem services are not included as high priority. A stratified approach may ensure that all native ecosystems are represented in the expanding PA network. Second, we are using models of current imperiled species distributions to infer the general patterns of protections, some of which may shift with global climate change (see 24). Future local, regional and continental scale analyses can help inform which areas need long-term protections. Finally, the GAP classification definitions may not account for substantive protections observed on the ground. For example, Department of Defense installations represent 20 million acres of GAP 4 land (some with high imperiled species diversity; 25) and have Integrated Natural Resource Management Plans that address some biodiversity concerns. There is also a great need for incorporation of Tribal knowledge as Tribal areas have some of the lowest rates of habitat modification, yet much of the over 56 million acres held in trust by the Bureau of Indian Affairs are not well documented in PADUS (26). These and similar situations highlight the nuances of inferring protection status.

Achieving 30x30 to help protect biodiversity and address the climate crisis in the U.S. is feasible but will require partnerships with nongovernmental landowners and across levels of government. Our analysis recognizes that the approaches and policy tools for doing so will vary considerably throughout the country. The key to operationalizing 30x30 will be planning beyond the numerical target for a protected areas network that can be established in a way that ensures a long-term commitment to biodiversity and climate. By doing so, the U.S. can continue to lead the way globally in protecting nature for its own sake and for our health and well-being.

## Materials and Methods

### Data Layers and Sources

Data layers were chosen that represent a recent baseline in the main objectives of the 30x30 framework: imperiled species biodiversity metrics represent species in greatest need of conservation and generally exhibit spatial patterns consistent with other measures of biodiversity (10) and ecosystem carbon stores represent areas that, if disturbed, could contribute additional carbon to the atmosphere. Terrestrial imperiled species richness and rarity-weighted richness from the Map of Biodiversity Importance project database (27) are based on habitat suitability models for 2,216 species and 11 taxa. Species richness is useful for understanding where ranges of the most species overlap a single location, but could be a poor indicator of diversity because it does not account for complementarity among sites. Alternatively, rarity-weighted richness – where species are assigned a value inversely proportional to the size of their range - may be a better method for maximizing the representation of species (28). Therefore, we take into account the number of species, proportional abundance of species presence, and the occurrence of rare species of conservation priority. Total ecosystem carbon is based on a map of global above- and below-ground carbon stored in biomass and soil in tonnes C per hectare (11).

Data on protected areas are from the PADUS v2.1 database (13). The layer was flattened prior to spatial overlay analysis to avoid overlaps in land units. This was done so in a way to give preference to the lowest GAP code present in any single location (i.e., greatest protection). U.S. terrestrial boundaries reflect all states and territories from the U.S. Geological Survey.

### Spatial Analyses

We used spatial overlay analysis to describe the extent to which the PA network covers U.S. lands and seas as well as areas of high imperiled species biodiversity and ecosystem carbon (see SI for marine results). Areas considered hotspots of biodiversity or carbon richness were in the 90^th^ percentile of the distribution of values for the respective metric. We used the associated Gap Analysis Program (GAP) codes, which are specific to the management intent to conserve biodiversity. GAP 1 and 2 areas are managed in ways typically consistent with conservation. GAP 3 areas are governed under multiple-use mandates and GAP 4 areas lack any conservation mandates (see Panel 1). Additionally, we summarized statistics by lands management (e.g., manager type and name attribute) and state or territory.

Imperiled species richness and carbon layers were also compared to data representing two major risks faced by unprotected species and landscapes: land conversion and climate change. Projections of future U.S. land cover in 2050 given the continuation of land-use trends from 1992-1997 were used to estimate the probability (scale of 0-1) of conversion of each location (1km^2^) to non-natural land uses (i.e., urban, crop or pasture; 29). Forward climate velocity, the speed (in km/year) at which species must migrate to maintain constant climate conditions, is a metric used to evaluate the exposure of places to climate change (30). The two risk variables were also combined in overlay analyses to evaluate unprotected hotspots of biodiversity and carbon (i.e., locations in the 90^th^ percentile of the distribution of values that did not overlap with GAP 1 and 2 areas). We used ArcPro v. 2.7.1 (ESRI, USA) to produce maps and run analyses. Maps use the Albers Equal Area Conic, Alaska Albers, and Old Hawaiian UTM Zone 4 projections.

## Acknowledgments

We thank T Niederman and A Carter for their thoughtful review of this manuscript. We also thank USGS for making PADUS, C. Soto-Navarro for making above- and below-ground carbon, and NatureServe and Esri for making the MoBI layers available to streamline general analyses.

## Author contributions

The authors contributed to this work in the following ways:

Conceptualization: LMD, JWM
Methodology: LMD, JWM
Investigation: LMD
Visualization: LMD, JWM
Supervision: LMD, JWM
Writing—original draft: LMD, JWM
Writing—review & editing: LMD, JWM

## Data and materials availability

All data used in the analyses are publicly available through the cited resources.

---

**Panel 1.**
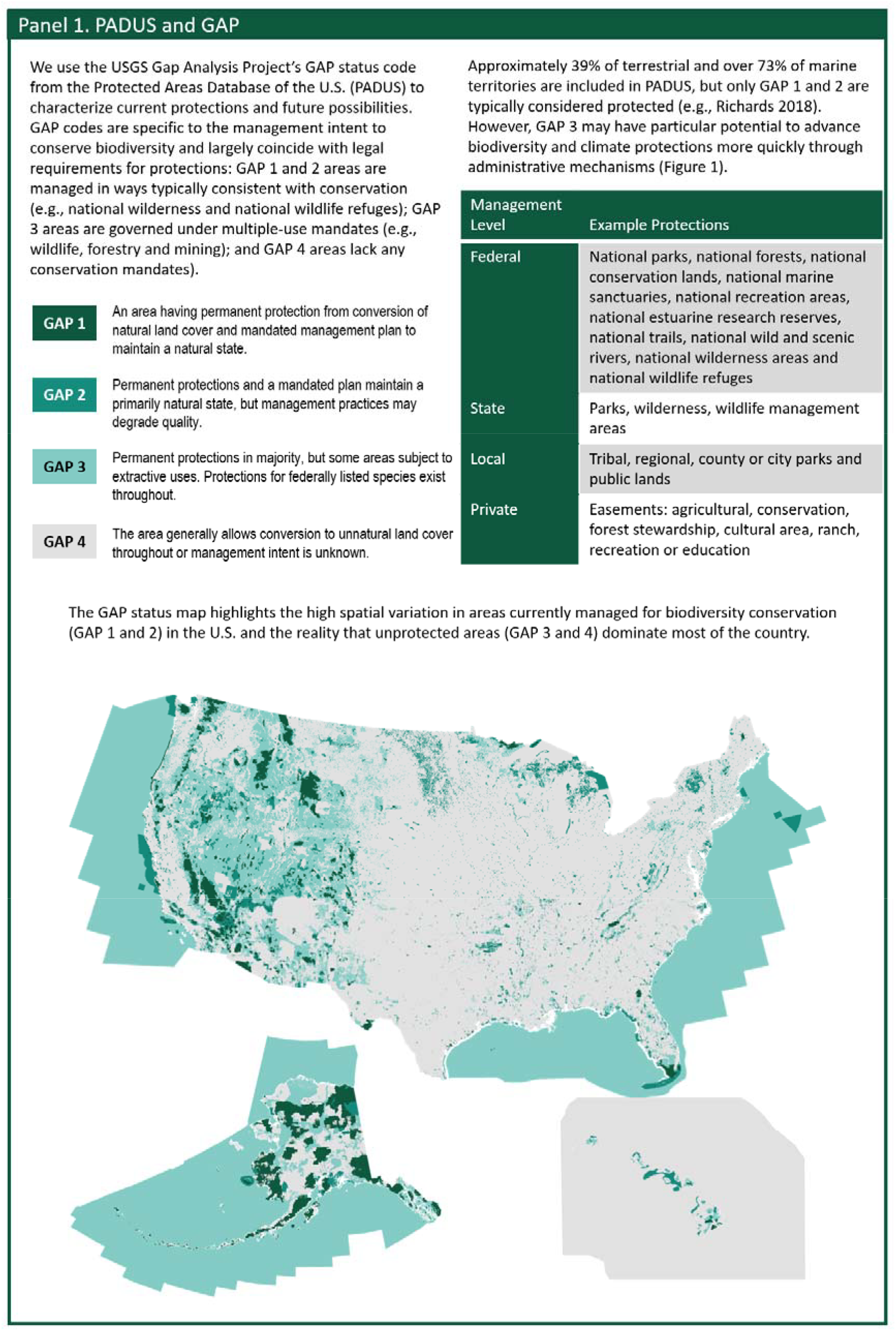
The Protected Areas Database of the US GAP codes explained.

## References

1. IPBES Secretariat, “Global assessment report on biodiversity and ecosystem services of the Intergovernmental Science-Policy Platform on Biodiversity and Ecosystem Services”, (Bonn, Germany, 2019).

2. Secretariat of the Convention on Biological Diversity, “COP-10 Decision X/2. Secretariat of the convention on biological diversity” (Convention on Biological Diversity, 2010)

3. R.P. Powers, W. Jetz, Global habitat loss and extinction risk of terrestrial vertebrates under future land-use-change scenarios. Nat. Clim. Chang. 9, 323–329 (2019).

4. E. Dinerstein, C. Vynne, E. Sala, A.R. Joshi, S. Fernando, T.E. Lovejoy, J. Mayorga, D. Olson, G.P. Asner, J.E.M. Baillie, N.D. Burgess, K. Burkart, R.F. Noss, Y.P. Zhang, A. Baccini, T. Birch, N. Hahn, L.N. Joppa, E. Wikramanayake, A global deal for nature: guiding principles, milestones, and targets. Sci. Adv. 5, eaaw2869 (2019).

5. Secretariat of the Convention on Biological Diversity, “Zero draft of the Post-2020 Global Biodiversity Framework” (Convention on Biological Diversity, 2020).

6. J.R. Biden, “Executive Order on Tackling the Climate Crisis at Home and Abroad (2021: https://www.whitehouse.gov/briefing-room/presidential-actions/2021/01/27/executive-order-on-tackling-the-climate-crisis-at-home-and-abroad/).

7. G. Newsom, “Exec Order N-82-20” (2020: https://www.gov.ca.gov/wp-content/uploads/2020/10/10.07.2020-EO-N-82-20-signed.pdf).

8. P. Kulberg, E. Di Minin, A. Moilanen, Using key biodiversity areas to guide effective expansion of the global protected areas network. Glob. Ecol. Conserv. 20, e00768 (2019).

9. R.T. Belote, M.S. Dietz, C.N. Jenkins, P.S. McKinley, G.H. Irwin, T.J. Fullman, J.C. Leppi, G.H. Aplet, Wild, connected, and diverse: building a more resilient system of protected areas. Ecol. Appl. 27, 1050–1056 (2017).

10. C.N. Jenkins, K.S. Van Houtan, S.L. Pimm SL, J.O. Sexton, US protected lands mismatch biodiversity priorities. PNAS 112, 5081–5086 (2015).

11. C. Soto-Navarro, C. Ravilious, A. Arnell, X. de Lamo, M. Harfoot, S.L.L. Hill, O.R. Wearn, M. Santoro, A. Bouvet, S. Mermoz, T. Le Toan, J. Xia, S. Liu, W. Yuan, S.A. Spawn, H.K. Gibbs, S. Ferrier, T. Harwood, R. Alkemade, A.M. Schipper, G. Schmidt-Traub, B. Strassburg, L. Miles, N.D. Burgess, V. Kapos, Mapping co-benefits for carbon storage and biodiversity to inform conservation policy and action. Phil. Trans. Royal Soc. B 375, doi: 10.1098/rstb.2019.0128 (2020).

12. D. Stralberg, C. Carroll, S.E. Nielsen, Toward a climate-informed North American protected areas network: Incorporating climate-change refugia and corridors in conservation planning. Conserv. Lett. 13, ee12712 (2020).

13. U.S. Geological Survey (USGS) Gap Analysis Project (GAP), Protected Areas Database of the United States (PAD-US) (2020; doi: 10.5066/P955KPLE).

14. J.J. Lawler, A.S. Ruesch, J.D. Olden, B.H. McRae, Projected climate-driven faunal movement routes. Ecol. Letters 16, 1014–1022 (2013).

15. Eichenwald

16. M.D. Barnes, L. Glew, C. Wyborn C, I.D. Craigie, Prevent perverse outcomes from global protected area policy. Nat. Ecol. Evol. 2, 759–762 (2018).

17. G. Baldi, M. Texeira, O.A. Martin OA, H.R. Grau, E.G. Jobbágy, Opportunities drive the global distribution of protected areas. PeerJ 5, e2989 (2017).

18. US Departments of the Interior, Agriculture, Commerce, Council on Environmental Quality, “Conserving and Restoring American the Beautiful” (https://www.doi.gov/sites/doi.gov/files/report-conserving-and-restoring-america-the-beautiful-2021.pdf, 2021).

19. J. Michalak, J. Lerner, “Linking conservation and land use planning: using the State Wildlife Action Plans to protect wildlife from urbanization” presented at the Transportation Land Use, Planning, and Air Quality Congress, American Society of Civil Engineers, Orlando, FL, 2007.

20. U.S. 116th Congress, House Resolution 3742 ‘Recovering America’s Wildlife Act’ (2019; https://www.congress.gov/bill/116th-congress/house-bill/3742)

21. B.B.N. Strassburg, A. Iribarrem, H.L. Beyer, C.L. Cordeiro, R. Crouzeilles, C.C. Jakovac, A.B. Junqueira, E. Lacerda, A.E. Latawiec, A. Balmford, T.M. Brooks, S.H.M. Butchart, R.L. Chazdin, K.H. Erb, P. Brancalion, G. Buchanan, D. Cooper, S. Diaz, P.F. Donald, V. Kapos, D. Leclere, L. Miles, M. Obersteiner, C. Plutzar, C.A. de M. Scaramuzza, F.R. Scarano, P. Visconti, Global priority areas for ecosystem restoration. Nature 586, 1–6 (2020).

22. E. Wood, A. Harsant, M. Dallimer, A.C. de Chavez, R.R.C. McEachan, C. Hassall, Not All Green Space Is Created Equal: Biodiversity Predicts Psychological Restorative Benefits From Urban Green Space. Front. Psychol. 9, doi: 10.3389/fpsyg.2018.02320 (2018).

23. E.N. Ussery, L. Yngve, D. Merriam, G. Whitfield, S. Foster, A. Wendel, T. Boehmer, The national public health tracking network access to parks indicator: a national county-level measure of park proximity. J. Park Recreat. Admi. 34, 52–63 (2016).

24. P.R. Elsen, W.B. Monahan, E.R. Dougherty, A.M. Merenlender, Keeping pace with climate change in global terrestrial protected areas. Sci. Adv. 6, eaay0814 (2020).

25. B.A. Stein, C. Scott, N. Benton, Federal lands and endangered species: the role of military and other federal lands in sustaining biodiversity. BioSci. 58, 339–347 (2008).

26. C. Vincent, L.A. Hanson, C. Argueta, “Federal Land Ownership: Overview and Data” (CRS Report R42346, 2020).

27. NatureServe Network, Map of Biodiversity Importance data collection (2020; https://livingatlas.arcgis.com/en/browse/#d=2&q=mobi).

28. F.D. Albuquerque, A. Gregory, The geography of hotspots of rarity-weighted richness of birds and their coverage by Natura 2000. PLoS One 12, e0174179 (2017).

29. V.C. Radeloff, E. Nelson, A.J. Plantinga, D.J. Lewis, D. Helmers, J.J Lawler, J.C. Withey, F. Beaudry, S. Martinuzzi, V. Bustic, E. Lonsdork, D. White, S. Polasky, Economic-based projections of future land use in the conterminous United States under alternative policy scenarios. Ecol. Appl. 22, 1036–1049 (2012).

30. C. Carroll, J.J. Lawler, D.R. Roberts, A. Hamann, Biotic and climatic velocity identify contrasting areas of vulnerability to climate change. PLOS ONE 10, e0140486 (2015).

31. United Nations Environment Programme –World Conservation Monitoring Center, “United Nations List of Protected Areas” (2018).

