## Supplemental Materials for "Identifying key federal, state and private lands strategies for achieving 30x30 in the US"

Supplementary Text

Marine Analyses

We conducted a parallel analysis for U.S. marine areas where marine boundaries reflect territorial waters (generally, three nautical miles from shore with some exceptions) and the exclusive economic zone (EEZ) for states and national analyses, respectively. This is to reflect the differences in how federal and state policies define waters included in the calculation of 30%: federal resolutions report on all PAs in the EEZ and the California state Executive Order count only state managed PAs within the state’s territorial waters. Marine imperiled species richness data are from the International Union for the Conservation of Nature and represent 254 species ranges and 9 taxa (10). Relative to international efforts, the U.S. ranks highly in marine protected area (PA) coverage (31).

Data show that 26% of U.S. seas that make up the exclusive economic zone are protected at levels consistent with the biodiversity and climate goals of 30x30 proposals (i.e., GAP 1 and 2). Up to 73.7% of marine habitats in the U.S. would be protected if regulatory changes to GAP 3 PAs emphasized biodiversity protection over other uses. Protections vary widely yet systematically across federal agencies, which has significant implications for how federal agencies will contribute to achieving 30x30. While there remains significant potential for achieving a national 30x30 numerically, high spatial variability in current PA designations will mean considerable heterogeneity in how numerical goals can be met at state levels.

Just under half of all U.S. states have marine PAs, which are located on coastlines and in the Great Lakes. There is high variability among states and regions in GAP 1 and 2 PA coverage (Table S2); spatial patterns of federally managed PAs indicate significant gaps in coastal protections including the northeast and the Gulf of Mexico where industrial fisheries are particularly active. If taking into account all PAs in respective territorial waters, four states have already achieved 30% protections within state territorial waters (Table S2). However, once constraining the PA network to state managed seas, all states fall short of the 30% target, with only a few (Hawaii, Florida) over half way. Including GAP 3 seas does not change this outlook.

Overall, 97% of seas that fall into GAP 3 classifications are federally managed leaving little room for improvement on the state level. Alaska, Massachusetts and North Carolina are interesting exceptions to this rule: if protections were to be strengthened on stage-managed GAP 3 PAs, 51%, 93%, and 41% of the marine zone would serve biodiversity conservation efforts in these states, respectively.

Table S1. Percentage of multiple use lands (GAP 3) by manager type and state jurisdiction.

| **% GAP 3 managed by** | | | | |  | **% GAP 3 managed by** | | | | |
| --- | --- | --- | --- | --- | --- | --- | --- | --- | --- | --- |
|  | **Federal** | **State** | **Local** | **Other** |  |  | **Federal** | **State** | **Local** | **Other** |
| Alabama | 72.67 | 17.74 | 9.11 | 0.48 |  | Montana | 79.02 | 20.65 | 0.05 | 0.28 |
| Alaska | 91.04 | 8.96 | 0.00 | 0.00 |  | Nebraska | 89.67 | 0.46 | 0.04 | 9.82 |
| Arizona | 96.49 | 2.98 | 0.34 | 0.19 |  | Nevada | 99.82 | 0.18 | 0.00 | 0.00 |
| Arkansas | 98.11 | 1.89 | 0.00 | 0.00 |  | New Hampshire | 57.19 | 15.42 | 7.41 | 19.98 |
| California | 93.87 | 2.01 | 3.77 | 0.35 |  | New Jersey | 7.25 | 0.58 | 20.98 | 71.18 |
| Colorado | 86.64 | 12.47 | 0.10 | 0.80 |  | New Mexico | 99.38 | 0.40 | 0.03 | 0.19 |
| Connecticut | 3.22 | 88.12 | 2.70 | 5.96 |  | New York | 2.47 | 94.87 | 1.88 | 0.78 |
| Delaware | 0.13 | 63.03 | 20.74 | 16.10 |  | North Carolina | 57.13 | 37.22 | 4.47 | 1.19 |
| Florida | 30.99 | 24.07 | 25.69 | 19.25 |  | North Dakota | 64.11 | 35.89 | 0.00 | 0.00 |
| Georgia | 94.63 | 5.30 | 0.06 | 0.00 |  | Ohio | 28.90 | 54.98 | 9.79 | 6.33 |
| Hawaii | 38.00 | 49.67 | 0.36 | 11.96 |  | Oklahoma | 70.10 | 22.72 | 3.38 | 3.80 |
| Idaho | 91.41 | 8.58 | 0.00 | 0.01 |  | Oregon | 94.30 | 5.25 | 0.44 | 0.01 |
| Illinois | 8.43 | 87.36 | 4.21 | 0.00 |  | Pennsylvania | 14.26 | 85.58 | 0.01 | 0.15 |
| Indiana | 51.11 | 48.18 | 0.40 | 0.31 |  | Rhode Island | 0.00 | 12.80 | 74.50 | 12.70 |
| Iowa | 8.32 | 91.27 | 0.03 | 0.38 |  | South Carolina | 79.65 | 17.61 | 0.33 | 2.42 |
| Kansas | 85.53 | 13.54 | 0.93 | 0.00 |  | South Dakota | 75.67 | 24.33 | 0.00 | 0.00 |
| Kentucky | 86.04 | 8.40 | 1.06 | 4.50 |  | Tennessee | 75.03 | 4.76 | 16.50 | 3.71 |
| Louisiana | 4.03 | 75.18 | 0.36 | 20.43 |  | Texas | 34.29 | 17.14 | 47.84 | 0.73 |
| Maine | 5.03 | 46.76 | 1.97 | 46.24 |  | Utah | 88.89 | 11.11 | 0.00 | 0.00 |
| Maryland | 18.83 | 79.94 | 0.00 | 1.23 |  | Vermont | 47.98 | 44.14 | 6.02 | 1.87 |
| Massachusetts | 0.83 | 61.91 | 30.15 | 7.12 |  | Virginia | 90.52 | 6.99 | 1.65 | 0.84 |
| Michigan | 0.02 | 99.96 | 0.00 | 0.02 |  | Washington | 71.11 | 28.57 | 0.26 | 0.06 |
| Minnesota | 26.71 | 73.28 | 0.00 | 0.00 |  | West Virginia | 91.58 | 8.17 | 0.24 | 0.01 |
| Mississippi | 92.54 | 2.31 | 0.28 | 4.87 |  | Wisconsin | 0.84 | 11.34 | 86.54 | 1.28 |
| Missouri | 99.95 | 0.00 | 0.00 | 0.05 |  | Wyoming | 85.91 | 13.41 | 0.00 | 0.68 |

Table S2. Marine protected area coverage by state, broken down by all vs. state-managed and by GAP status code. While some state marine zones (three nautical mile buffer, plus wider buffers for Texas, Puerto Rico, and parts of Florida) go beyond the 30% target, very few PAs are managed by states.

|  | **All (% Cover)** | | **State Managed  (% Cover)** | |
| --- | --- | --- | --- | --- |
|  | **GAP 1&2** | **GAP 1-3** | **GAP 1&2** | **GAP 1-3** |
| Alabama | 0.75 | 1.21 | 0.68 | 0.68 |
| Alaska | 7.05 | 87.33 | 0.81 | 50.87 |
| California | 43.00 | 53.63 | 13.10 | 13.10 |
| Connecticut | 0.00 | 99.37 | 0.00 | 0.00 |
| Delaware | 0.86 | 99.24 | 0.85 | 0.85 |
| Florida | 52.92 | 67.27 | 26.38 | 27.45 |
| Great Lakes | 8.50 | 8.51 | 2.74 | 2.75 |
| Georgia | 1.83 | 84.69 | 0.75 | 0.75 |
| Hawaii | 35.85 | 43.32 | 16.03 | 23.50 |
| Louisiana | 4.54 | 7.77 | 3.99 | 7.01 |
| Maine | 0.40 | 97.51 | 0.00 | 0.09 |
| Maryland | 0.76 | 98.70 | 0.06 | 11.54 |
| Massachusetts | 1.46 | 99.39 | 0.00 | 93.04 |
| Mississippi | 14.23 | 14.38 | 0.92 | 0.92 |
| New Hampshire | 0.24 | 99.13 | 0.25 | 1.29 |
| New Jersey | 5.16 | 99.29 | 4.77 | 4.82 |
| New York | 1.19 | 98.91 | 0.03 | 0.03 |
| North Carolina | 1.31 | 97.10 | 0.46 | 41.40 |
| Oregon | 10.18 | 10.19 | 9.82 | 9.82 |
| Rhode Island | 0.69 | 99.28 | 0.69 | 4.94 |
| South Carolina | 8.49 | 89.13 | 4.73 | 4.77 |
| Texas | 9.87 | 9.87 | 5.59 | 5.59 |
| Virginia | 1.04 | 98.23 | 0.02 | 17.18 |
| Washington | 45.56 | 45.57 | 0.07 | 0.07 |
